# Toward a global chemogeography of dissolved organic matter: a molecular trait dataset across Earth systems

**DOI:** 10.64898/2026.09.22.753682

**Authors:** Lei Han, Kang Song, James Stegen, Ang Hu

## Abstract

Dissolved organic matter (DOM) is a central component of biogeochemical cycling, but molecular observations generated by ultrahigh-resolution mass spectrometry remain fragmented across studies, limiting quantitative comparisons among ecosystems. Here, we compile and harmonize a global dataset of DOM molecular traits comprising 3,999 samples from 337 studies published between 2005 and 2026. The dataset includes eight intensity-weighted molecular traits: hydrogen-to-carbon (H/C), oxygen-to-carbon (O/C), nitrogen-to-carbon (N/C), and sulfur-to-carbon (S/C) ratios; molecular mass; double-bond equivalents (DBE); modified aromaticity index (AI_mod_); and nominal oxidation state of carbon (NOSC). Samples span aquatic, terrestrial, atmospheric, plant-associated, petroleum-related, and other environments, and are linked, where available, to geographic, physicochemical, nutrient, and climatic metadata. Molecular data were harmonized using common definitions and restricted to negative-mode electrospray-ionization FT-ICR MS to improve cross-study comparability. This resource enables cross-system comparisons of DOM molecular traits and supports analyses of environmental gradients, geographic coverage, and global DOM chemogeography.

**Highlights:**

1. A global dataset of DOM molecular properties was compiled from 3,999 samples reported in 337 publications.
2. The dataset integrates eight sample-level molecular traits: H/C, O/C, N/C, S/C, mass, DBE, AI_mod_, and NOSC.
3. Samples span six broad Earth-system categories and diverse habitats across land- to-ocean and other environmental settings.
4. Most records are linked to geographic, physicochemical, nutrient, and/or climatic metadata.
5. The dataset supports analyses of DOM chemogeography, habitat-associated variation, environmental drivers, and cross-system convergence.

## Introduction

Dissolved organic matter (DOM) constitutes one of the largest and most activity pools of organic carbon across Earth systems and plays key roles in carbon transport, nutrient cycling, microbial metabolism, and biogeochemistry (Battin et al., 2009; Dittmar et al., 2021; Jiao et al., 2010). Its ecological and biogeochemical functions are closely linked to molecular traits that vary markedly across terrestrial, freshwater, marine, atmospheric, and other environments (Arrieta et al., 2015; D’Andrilli et al., 2022, 2019; Hu et al., 2025a). Over the past two decades, Fourier transform ion cyclotron resonance mass spectrometry (FT-ICR MS) has greatly advanced the molecular-level characterization of DOM by resolving thousands of molecular formulae within individual samples (Qi et al., 2022; Sleighter et al., 2010; Sleighter and Hatcher, 2007). Commonly reported molecular traits include H/C, O/C, DBE, AI_mod_, mass, and NOSC, which describe complementary dimensions of elemental composition, unsaturation, aromaticity, molecular size, and carbon oxidation state (Boye et al., 2017; Dwinandha et al., 2023; LaRowe and Van Cappellen, 2011). The van Krevelen diagram, which organizes molecular formulae according to H/C and O/C ratios, has become a widely used framework for comparing DOM composition among samples and systems (Kim et al., 2003; Laszakovits and MacKay, 2021; Wu et al., 2004). Together, these analytical advances have enabled increasingly broad comparisons of DOM composition across environmental gradients (Freeman et al., 2024; Hertkorn et al., 2016; Herzsprung et al., 2020).

Advances in computational processing and molecular formula annotation have enabled increasingly systematic characterization of DOM molecular composition across broad environmental gradients and at expanding dataset scales (Meng et al., 2025; Seitzinger et al., 2005). However, these observations remain dispersed among individual studies, and differences in sampling design, analytical platforms, ionization modes, data processing, and reporting practices hinder direct synthesis (Fu et al., 2025; Kellerman et al., 2014; Koch et al., 2007; Shen and Benner, 2026). Many studies have also focused on ecosystem-specific or regional gradients (Chen et al., 2022; Melendez-Perez et al., 2016), limiting standardized comparisons at the global scale. A harmonized dataset that links DOM molecular traits with geographic, climatic, and physicochemical information would provide a common basis for investigating DOM chemogeography, evaluating habitat-level similarities and differences, and examining environmental associations (Fu et al., 2019; Stegen and Goldman, 2018; Wang et al., 2015; Yi et al., 2025). Such a resource would also facilitate integration with molecular observations generated by emerging chromatographic and high-resolution mass spectrometric approaches.

To address these limitations, we assembled and harmonized a global dataset of DOM molecular traits from published FT-ICR MS studies across Earth systems. The dataset links standardized sample-level molecular traits with habitat classifications and available geographic, climatic, physicochemical, and nutrient metadata. It is intended to support reproducible cross-system comparisons of DOM composition and hypothesis testing on how source characteristics and environmental filtering are associated with DOM molecular composition at broad spatial scales.

## Methods

### Data collection

We systematically compiled peer-reviewed studies published before April 2026 that characterized DOM molecular composition using FT-ICR MS. Publications were identified through the Web of Science (Core Collection; http://www.webofknowledge.com) and Google Scholar (http://scholar.google.com) using the search terms “organic matter AND FT-ICR MS AND van Krevelen” (Han et al., 2024; Hu et al., 2025a). Initial screening of abstracts and methods yielded 1,230 potentially relevant full-text articles. We then assessed the full texts for eligibility and retained 3,999 samples from 337 publications according to the following criteria:

1. Studies were conducted in natural or engineered environments; samples derived from experimental manipulations were excluded. This restriction was applied to focus the compilation on observational characterizations of DOM in environmental settings.
2. Publications either directly reported sample-level DOM molecular traits or provided molecular-formula data from which these traits could be calculated. For consistency across studies, all sample-level traits were defined as intensity-weighted averages across all assigned molecular formulae within each sample. When the traits were not reported directly, they were recalculated from available formula-level data using this common definition. Specifically, each formula-specific molecular traits was multiplied by its relative peak intensity, summed across all detected formulae, and normalized by total signal intensity (Roth et al., 2019).
3. Only data generated by FT-ICR MS were retained, and the dataset was further restricted to measurements obtained using negative-mode electrospray ionization (ESI-) to reduce analytical heterogeneity among studies (Hawkes et al., 2020). Results obtained using other high-resolution mass spectrometric platforms, such as Orbitrap MS, were excluded. Metadata describing sample pretreatment, including solid-phase extraction (SPE) approaches, were recorded whenever available because SPE-based enrichment is widely used before FT-ICR MS analysis of DOM to improve recovery and ionization efficiency.

### Environmental and climatic data

To complement the DOM molecular traits, we compiled physicochemical and climatic covariates for each sample from the original publications and the WorldClim database. Three geographic variables and 15 environmental variables were collected from the source studies, including latitude, longitude, altitude, temperature, pH, salinity, conductivity (EC), dissolved oxygen (DO), total organic carbon (TOC), total nitrogen (TN), total dissolved nitrogen (TDN), dissolved organic carbon (DOC), ammonia nitrogen (NH_4_^+^-N), nitrate nitrogen (NO_3_^-^-N), nitrite nitrogen (NO_2_^-^-N), orthophosphate (PO_4_^3-^-P), iron (Fe), and manganese (Mn).

Climatic conditions were further characterized using georeferenced sample coordinates and digital elevation data at a spatial resolution of 0.5^°^. For each sampling location, we extracted the 19 standard bioclimatic variables from WorldClim (Hijmans et al., 2005; Hu et al., 2025a): annual mean temperature (BIO1), mean diurnal range (BIO2), isothermality (BIO3), temperature seasonality (BIO4), maximum temperature of warmest month (BIO5), minimum temperature of coldest month (BIO6), temperature annual range (BIO7), mean temperature of wettest quarter (BIO8), mean temperature of driest quarter (BIO9), mean temperature of warmest quarter (BIO10), mean temperature of coldest quarter (BIO11), annual precipitation (BIO12), precipitation of wettest month (BIO13), precipitation of driest month (BIO14), precipitation seasonality (BIO15), precipitation of wettest quarter (BIO16), precipitation of driest quarter (BIO17), precipitation of warmest quarter (BIO18), and precipitation of coldest quarter (BIO19).

### Data records

The compiled dataset is provided as Supplementary Table S1.

### Data overview

The dataset comprises 3,999 samples collected between 2003 and 2026 across diverse Earth systems (Fig. 1). Five major categories, including waters, land, atmosphere, plant-associated, and petroleum-related environments, contained 2,696, 534, 178, 152, and 101 samples, respectively, with additional samples assigned to other environmental categories. The number of DOM samples analyzed by ultrahigh-resolution mass spectrometry increased sharply after 2015, reflecting the expanding application of FT-ICR MS in DOM research. Water samples represented 69.7% of the dataset and land samples 13.8%. Sampling locations spanned North and South America, Europe, Asia, Africa, Oceania, and polar regions, encompassing broad climatic and environmental gradients. Geographic coverage was nevertheless uneven, with records concentrated in North America, Europe, and East Asia and relatively sparse representation in Africa, central Asia, several open-ocean regions, and high-latitude environments. This spatial imbalance should be considered when the dataset is used for global inference.

**Figure 1.**
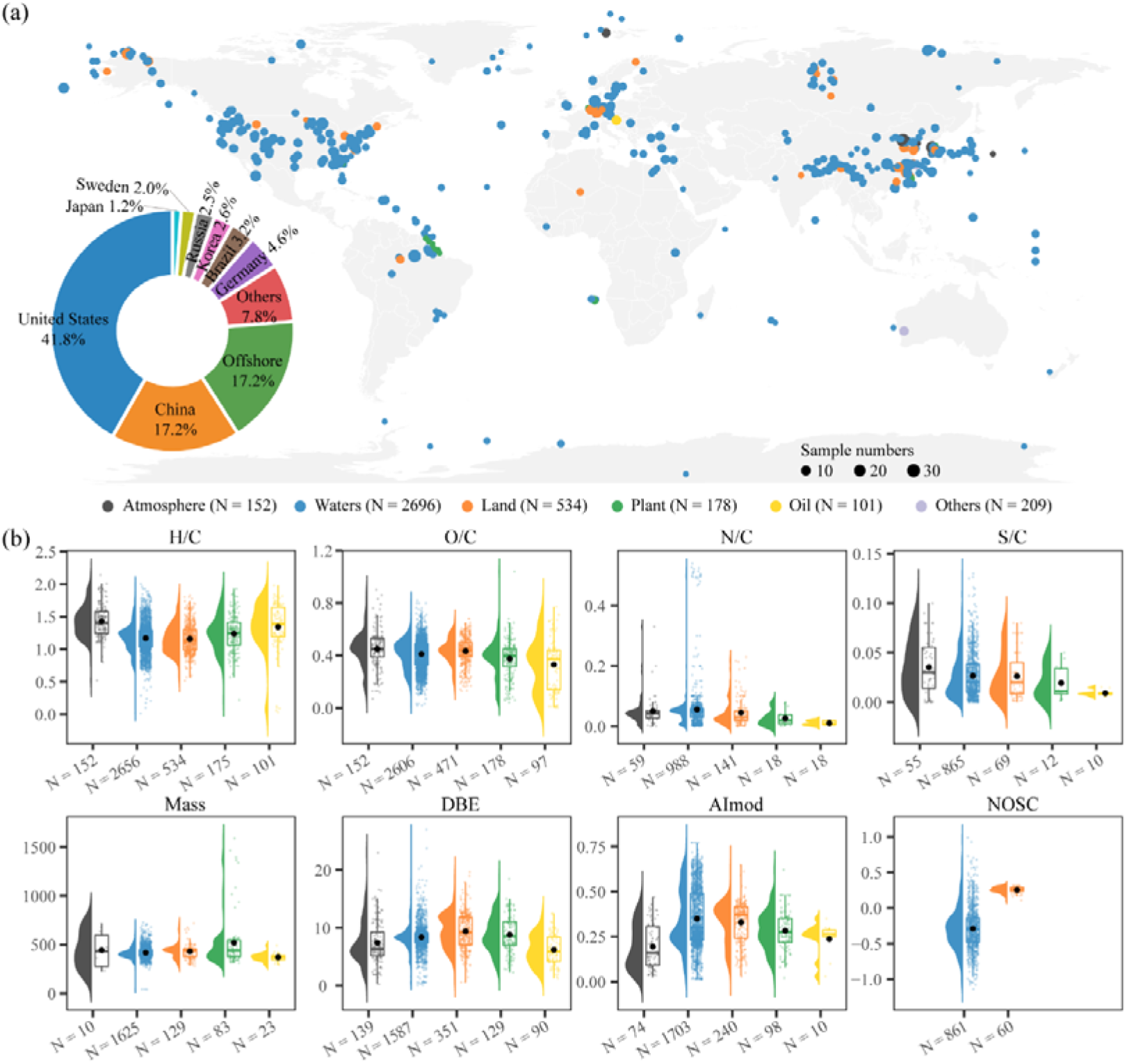
Global distribution of samples and variation in DOM molecular traits among Earth systems. (a) Global distribution of samples collected from six ecosystems. Point colors represented environmental categories, while point sizes indicated the number of samples at each sampling location. The inset donut chart showed the proportional contribution of samples from different countries. (b) Variation in eight molecular traits across Earth systems. Half-violin plots illustrated the overall data distributions, overlaid points displayed individual observations, and boxplots summarized the median and interquartile range, and the N represented the sample sizes and the black dots in the Boxplots represented the mean values.

DOM molecular traits varied substantially across the compiled records (Figs. 1, 2). Dataset availability was greatest for H/C and O/C ratios, with 3,616 and 3,504 observations, respectively, followed by DBE (2,296), AI_mod_ (2,125), molecular mass (1,870), N/C (1,224), S/C (1,011), and NOSC (921), primarily from aquatic and terrestrial systems. Trait distributions showed broad ranges and varying degrees of overlap among Earth systems (Fig. 1b). Specifically, H/C ratios were concentrated largely between approximately 1.0 and 1.6, with generally higher values in atmospheric and petroleum-associated samples than in water and land samples. This pattern is consistent with greater hydrogen saturation in some atmospheric, plant-derived, and petroleum-associated organic matter and relatively greater unsaturation in many aquatics and terrestrial DOM pools. Most O/C ratios occurred between approximately 0.3 and 0.6; atmospheric DOM tended toward higher values, whereas petroleum-related samples occupied the lower end of the range. In contrast, waters, land, and plant samples showed substantial overlap, suggesting that O/C alone provides limited separation among Earth systems. N/C and S/C were strongly concentrated near zero and were strongly right skewed, with a small number of comparatively high observations. Most samples contained relatively low intensity-weighted contributions of nitrogen- and sulfur-containing molecular formulae, while several samples exhibited markedly elevated values. Such like-outlier values emphasized the importance of retaining the continuous molecular information rather than classifying DOM solely according to vague CHO, CHON, or CHOS formula groups. Molecular mass was generally between 300 and 500 Da, although plant-associated DOM spanned a broader range and included values above 1,500 Da. DBE values were mostly between 5 and 12, with terrestrial and plant DOM exhibiting slightly higher median values than atmospheric and petroleum samples. AI_mod_ was lower in atmospheric DOM, with values around 0.2, and higher in aquatic and terrestrial DOM, with values of approximately 0.3-0.4. NOSC was predominantly negative in waters, with values centered near -0.3, but positive in land DOM, with values around 0.2-0.3. Molecular traits also varied among individual habitats within the major Earth systems (Fig. 2), providing an empirical quantitative basis for characterizing the biogeographic patterns of DOM chemistry across diverse global environments (He et al., 2023; Hu et al., 2025b).

**Figure 2.**
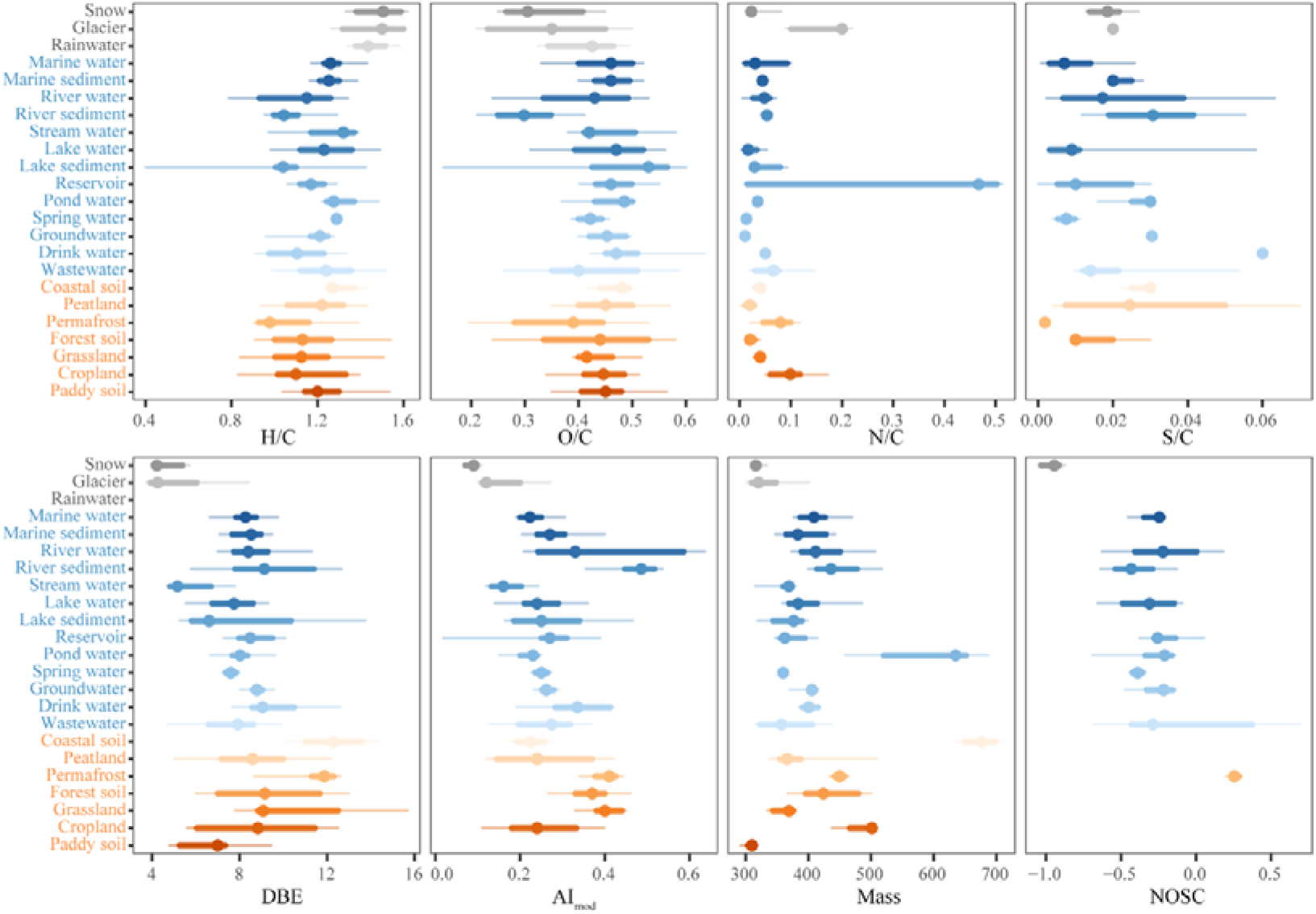
Comparisons of DOM molecular traits across water and land systems. Points represented the median values, and the thick and thin horizontal bars indicated the interquartile ranges of 25th-75th percentiles and 10th-90th percentiles, respectively. Water habitats are shown in grey and blue and land habitats in orange. Notes: the distinction between ‘river water’ and ‘stream water’ in the literature is often context-dependent and lacks a universally accepted definition. In our dataset, these categories follow the descriptions provided in the original studies. Because no strict boundary exists between streams and river—and many studies may classify similar systems differently—any observed differences between these two categories should be interpreted with caution. For analyses focused on broader environmental patterns, these two categories can be merged as ‘riverine water’ to reduce classification heterogeneity.

The database also spans broad geographic, climatic, and physicochemical gradients (Fig. 3). Sampling elevations ranged from locations near or below sea level to high-elevation environments above 6,000 m, with waters record spanning the widest altitudinal range. Mean annual temperature extended from below −10 °C in cold polar and high-altitude environments to above 30 °C in warm tropical and subtropical systems. Mean annual precipitation spanned strongly arid to humid environments, ranging from a few hundred millimeters to more than 3,000 mm yr^−1^. The 19 bioclimatic variables further capture variation in thermal seasonality, annual and diurnal temperature range, precipitation seasonality, and climatic extremes. This breadth is particularly valuable because global DOM composition may respond not only to long-term climatic means but also to seasonal and extreme environmental conditions that regulate hydrological connectivity, vegetation inputs, microbial and photochemical processing. Local physicochemical measurements were less consistently reported among studies: pH, salinity, and DOC were the most widely available, while nutrient and trace element factors, particularly TOC, TN, nitrite, phosphate, Fe, and Mn, were comparatively sparse. The number of available observations and the recorded ranges for all environmental variables are provided in metadata. Comparing these sampled ranges with the global distribution of environmental conditions could identify regions or environmental settings that are currently underrepresented in the dataset (Kong et al., 2025; Stegen et al., 2018). Such information may help guide future sampling efforts and improve the global representativeness of DOM datasets.

**Figure 3.**
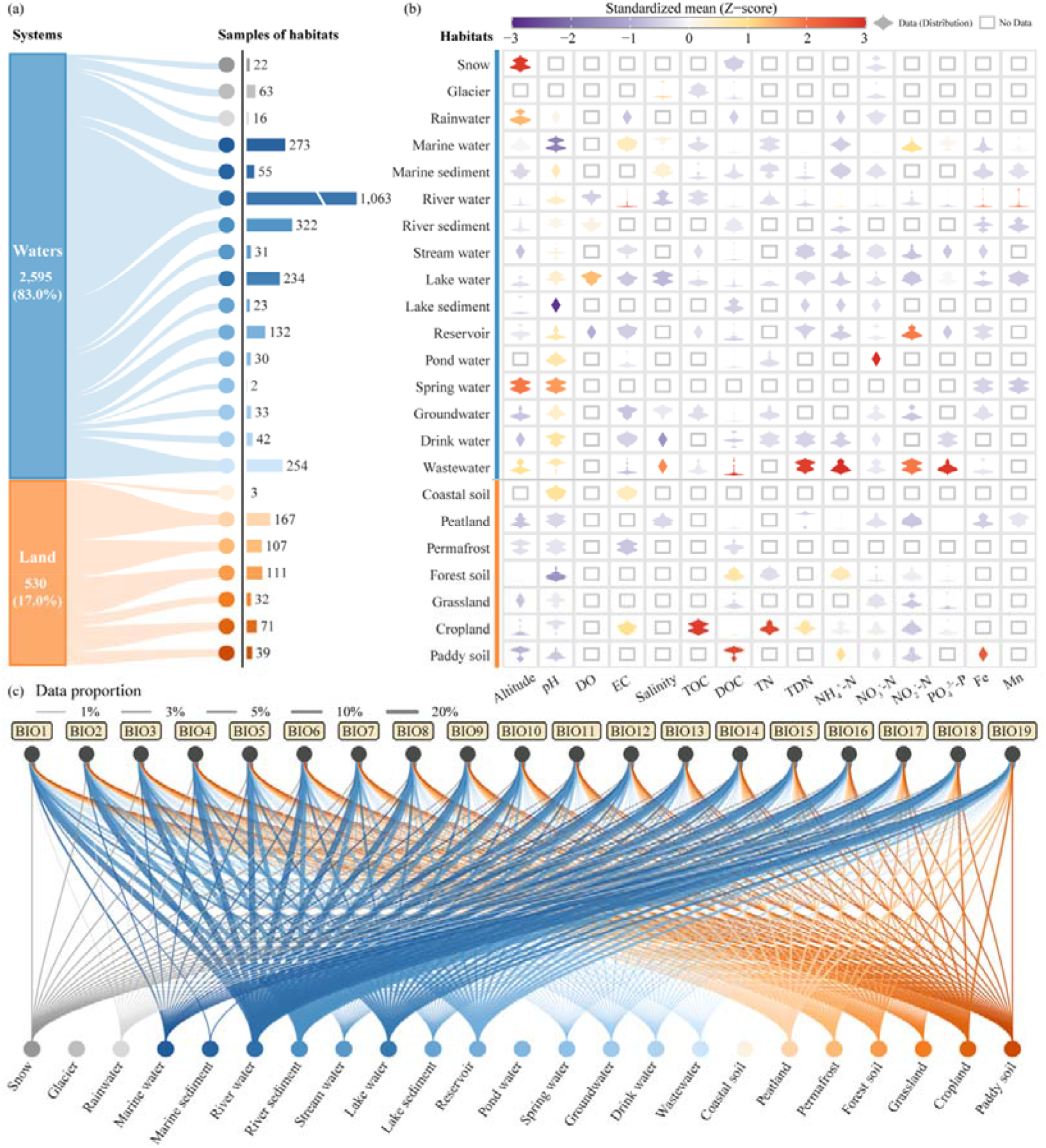
Environmental and climatic variables versus data coverage across water and land systems and habitats. (a) The left panel summarized the hierarchical structure, and samples were only showed two major systems (waters and land). The connecting ribbons link each system to its constituent habitats, whereas the adjacent horizontal bars indicated the number of observations available for each habitat. Habitats were exhibited in a predefined ecological order, with water habitats shown in blue and land habitat in orange. (b) The right panel depicted the distributions of environmental variables across the 23 major habitats using cell-wise density plots. For each environmental variable, habitat-level mean values were standardized across habitats as Z-scores, with colors indicating the relative deviation from the overall mean. The purple tones denoted below-average values, colors near the center of the scale indicated values close to the general mean, and orange-to-red tones displayed above-average values. The shape and width of each density glyph represent the within-habitat distribution of the corresponding environmental variable, while open gray squares indicated unavailable data. (c) The bottom panel visualized the coverage of the 19 bioclimatic variables (BIO1-BIO19) across all habitats by a bipartite network. The width of each connecting curve was proportional to the contribution of a given habitat to the total number of valid observations for the corresponding bioclimatic variable.

### Technological validation

The global DOM dataset was developed using a series of data-screening, harmonization, and quality-control procedures. Records that did not meet the predefined inclusion criteria or lacked sufficient methodological information for reliable interpretation were excluded. To reduce analytical heterogeneity among studies, datasets were standardized with respect to mass-spectrometric platform, ionization mode, and the calculation of sample-level molecular traits. For cross-system comparisons, only data generated by negative-mode ESI-FT-ICR MS were retained, and molecular traits were required to represent the complete set of assigned molecular formulae within each sample rather than selected molecular subsets. Variable names, units, and metadata definitions were harmonized across publications. Geographic coordinates, physicochemical measurements, and associated metadata were cross-checked against the original sources. Implausible or internally inconsistent records were re-examined and corrected when sufficient information was available; otherwise, they were designated as missing or excluded from the relevant analyses. Together, these procedures improved the consistency, traceability, and comparability of DOM molecular data compiled across diverse Earth systems and habitats.

### Usage Notes

The dataset provides a common framework for comparing DOM molecular traits among habitats and environmental settings that have often been studied separately (Fig. 4). Molecular traits can be examined together with habitat types, geographic location, climatic conditions, and available local environmental measurements. One potential use is to test the relative importance of source characteristics and environmental processing. Under stronger source control, DOM derived from similar biological or geological sources may retain comparable molecular signatures despite differences in local environmental conditions (Kalinski et al., 2026; Mostovaya et al., 2025; Wagner et al., 2015). In contrast, stronger environmental processing control may promote convergence among initially distinct DOM pools exposed to similar physicochemical, hydrological, photochemical, or microbial environments (Freeman et al., 2024; Song et al., 2022).

**Figure 4.**
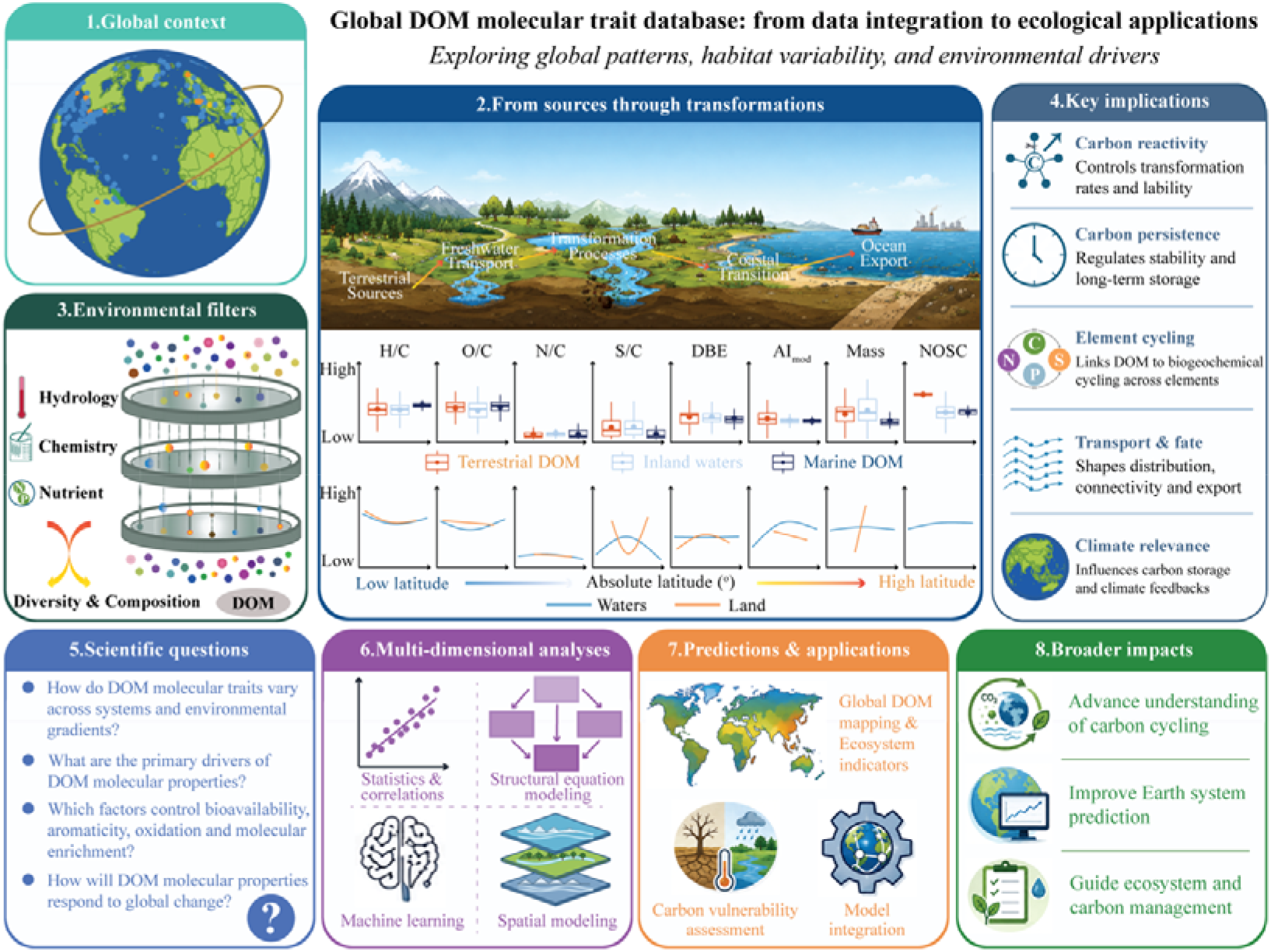
Conceptual framework linking environmental context, DOM molecular traits, and potential implications of the dataset. Global environmental context and local physicochemical conditions can influence DOM composition through source inputs, transport, and transformation processes. The integrated dataset enables cross-scale analyses of molecular trait variation, supports identification of key drivers, and provides a basis for predicting DOM dynamics, carbon cycling, and ecosystem responses under global change. Habitats were classified along the land-to-ocean continuum into three major ecosystem categories: Terrestrial DOM (e.g., permafrost, forest soil, grassland, cropland, paddy soil, peatland); Inland waters (e.g., river water, river sediment, stream water, lake water, lake sediment, reservoirs, pond water, spring water, groundwater); Ocean (e.g., marine water, marine sediment).

This source–processing framework is particularly relevant along the land-to-ocean continuum (Fettweis et al., 2026; Regnier et al., 2022; Ward et al., 2020). Organic matter originating from vegetation, soils, wetlands, atmospheric deposition, and cryosphere environments can be transported through groundwater, streams, rivers, lakes, reservoirs, estuaries, and ocean (Guo et al., 2025; Moran et al., 2022; Zhu et al., 2020). During transport, molecular composition can be modified by microbial degradation and production, photochemical transformation, sorption and desorption, mineral interactions, dilution, salinity change, and hydrological residence time (Chen et al., 2022; Díaz-Martínez et al., 2024; Freeman et al., 2024; Kang et al., 2023). A globally harmonized molecular dataset provides a framework for examining the relative importance of source imprints and environmental processing in shaping DOM molecular properties. By comparing molecular variation among habitats and environmental settings, the dataset can therefore be used to test whether habitat-associated molecular signatures persist across environmental gradients or converge under comparable conditions (Wang et al., 2024, 2023; Zhao et al., 2024). It may also support machine-learning, spatial-modeling, and ecosystem-scale applications to generate global DOM molecular mapping, assess ecosystem carbon vulnerability, and improve representation of organic matter cycling in Earth system models.

The dataset can also be integrated with emerging structure-resolved approaches, including LC–FT-ICR MS, LC–Orbitrap MS, and LC–MS/MS (Gao et al., 2026; Lu et al., 2015; Stumpf et al., 2025). Combining formula-level properties with chromatographic retention, fragmentation, isotopic, and transformation information could extend future DOM databases from descriptions of molecular composition toward more detailed characterization of molecular structure, reactivity, and fate (Matos et al., 2025; Spencer et al., 2026).

## Data availability

The compiled dataset is provided as Supplementary Table S1.

## Acknowledgments

This study was supported by National Natural Science Foundation of China (42622716). We acknowledge the use of artificial intelligence tools (ChatGPT) exclusively as a coding assistant and check English grammar. Authors reviewed and edited the content as needed and takes full responsibility for the content of the published article.

## Author contributions

AH conceived the review. LH synthesized and analyzed the data with the contributions of AH, KS, and JS. LH finished the first draft and finalized the manuscript with the contributions of all authors.

## Conflict of interests

The authors declare no conflicts of interest.

## Notes

### Competing Interest Statement

The authors have declared no competing interest.

